# Both lipopolysaccharide and lipoteichoic acids potently induce anomalous fibrin amyloid formation: assessment with novel Amytracker^™^ stains

**DOI:** 10.1101/143867

**Authors:** Etheresia Pretorius, Martin J Page, Lisa Hendricks, Nondumiso B Nkosi, Sven R Benson, Douglas B Kell

## Abstract

In recent work, we discovered that the presence of highly substoichiometric amounts (10^−8^ molar ratio) of lipopolysaccharide (LPS) from Gram-negative bacteria caused fibrinogen clotting to lead to the formation of an amyloid form of fibrin. We here show that the broadly equivalent lipoteichoic acids (LTAs) from two species of Gram-positive bacteria have similarly (if not more) potent effects. Using thioflavin T fluorescence to detect amyloid as before, the addition of low concentrations of free ferric ion is found to have similar effects. Luminescent conjugated oligothiophene dyes (LCOs), marketed under the trade name Amytracker^TM^, also stain classical amyloid structures. We here show that they too give very large fluorescence enhancements when clotting is initiated in the presence of the four amyloidogens (LPS, ferric ions and two LTA types). The staining patterns differ significantly as a function of both the amyloidogens and the dyes used to assess them, indicating clearly that the nature of the clots formed is different. This is also the case when clotting is measured viscometrically using thromboelastography. Overall, the data provide further evidence for an important role of bacterial cell wall products in the various coagulopathies that are observable in chronic, inflammatory diseases. The assays may have potential in both diagnostics and therapeutics.

## Introduction

In recent work, we have developed the idea that lipopolysaccharides (LPS) from the Gram-negative cell envelope can be shed from dormant bacteria or from continual bacteria entry into the blood e.g. via a leaky gut, and serve to contribute to the chronic inflammation characteristic of a considerable number of diseases [1–6]. Coupled to iron dysregulation [7], this leads to various coagulopathies [8] and changes in the morphology [9] both of erythrocytes and of the fibrin formed following blood clotting. A particularly striking finding was the fact that this ‘anomalous’ fibrin (sometimes referred to by us as ‘dense matter deposits’ (DMDs), e.g. [10, 11]) could be induced by tiny amounts of LPS amounting to 1 LPS molecule per 10^8^ molecules of fibrinogen [12]. This kind of sub-stoichiometric or autocatalytic activity was reminiscent of prion-like or β-amyloid behaviour. Indeed the ‘anomalous’ fibrin was found to be stainable by the amyloid-selective stain Thioflavin T (ThT), and hence amyloid in nature [12]. This provided a straightforward explanation for a number of elements of its biology, not least its resistance to degradation by the usual enzymes [8, 11].

This narrative could account for the role of Gram-negative bacteria, but not for that of Gram-positives, as these do not possess LPS. By contrast, their cell walls contain lipoteichoic acids (LTA), soluble peptidoglycan and muropeptides. There is a general feeling [13], especially since LTA has been properly purified [14], that LTA is just as capable as is LPS of providing an inflammatory stimulus to cells. While LPS mainly interacts with toll-like receptor 4 (TLR4) [2, 15–17], LTA stimulates target cells mainly by activating toll-like receptor 2 [13, 18–28], and with the glycolipid anchor of LTA playing a central role, analogous to the lipid A of LPS [29]. This is reasonable, as from the host’s point of a view an invading microorganism is simply undesirable, whatever its reaction to the Gram stain. Indeed, modulo a few detailed differences [30], and certainly for our present purposes, it seems that LTA is indeed broadly equivalent to LPS in terms of its ability to stimulate an innate immune response [31–33].

As well as the well-known ThT (e.g. [11, 34–50]), a considerable number of other fluorogenic stains have been shown to illuminate amyloids (e.g. [11, 51–67]). In particular, Nilsson and colleagues have developed a number of novel fluorescent amyloidogenic markers. Some of these are referred to as luminescent conjugated oligothiophenes (LCOs) [68–76] and are marketed as Amytracker™ 480 and 680 (derived, respectively, from HS163 and hFTAA HS169 in the literature [71, 74]), but proprietary structures that are not identical to them; (see Figure 1 for the chemical structures of HS163 and HS169). Although they too show strong selectivity for, and enhanced fluorescence when binding to, classical amyloid proteins, their staining properties clearly differ from those of ThT [68, 70, 77–79]. In some cases, their binding affects prion formation itself [80, 81] (and even ThT has anti-ageing properties at low concentrations [82]). It was thus of interest to assess these too as amyloid markers of the fibrin(ogen) blood clotting system.

**Figure 1:**
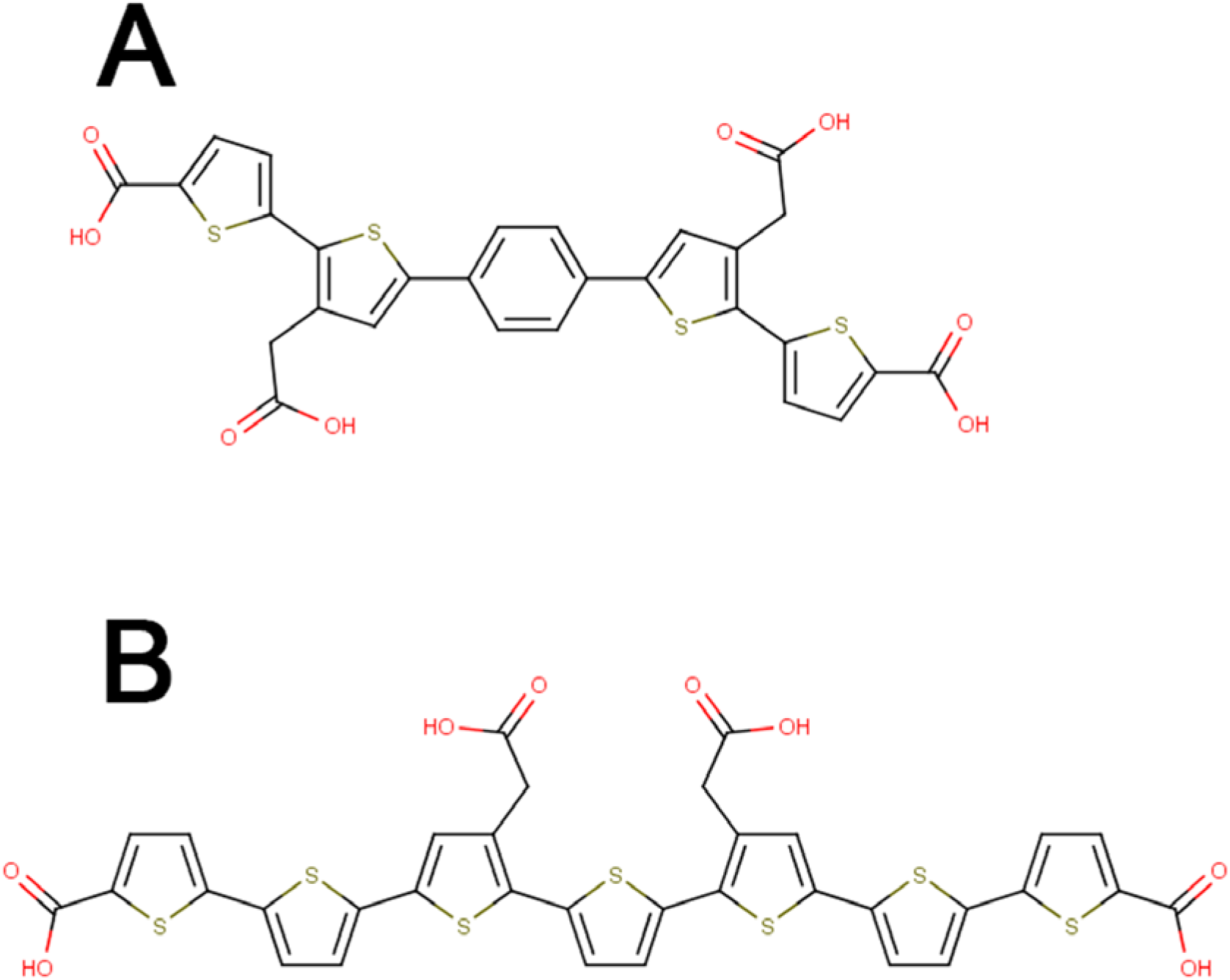
Chemical structures and SMILES [83] representation of **(A)** HS163 - SMILES (OC(=O)CC1=C(SC(=C1)C1=CC=C(C=C1)C1=CC(CC(O)=O)=C(S1)C1=CC=C(S1)C(O)=O) C1=CC=C(S1)C(O)=O) and **(B)** HS169 - SMILES O=C(O)c1ccc(s1)c2sc(cc2CC(=O)O)c3ccc(c4nsnc34)c5cc(CC(=O)O)c(s5)c6ccc(s6)C(=O)O (structures taken from [84]).

The question then arose as to whether LTA and unliganded iron (which is also almost always dysregulated during inflammation [7, 85, 86]), could display just as strong an ability to divert fibrinogen polymerisation from the normal to the amyloid form as could LPS. This was very much the case, and the present study shows the amyloid-forming nature of blood clots formed in plasma in the presence of low concentrations of iron, LPS from *E. coli*, and two LTAs from *Staphylococcus aureus* and *Streptococcus pyogenes.* These latter were chosen on the basis of coming from infectious_7_ Gram-positive organisms. A preprint has been lodged at bioRxiv [87].

## Materials and Methods

### Ethical statement

This study was approved by the Ethical Committee of the University of Pretoria (South Africa) and Stellenbosch University: ethics clearance number: 298/2016. We adhered strictly to the Declaration of Helsinki. A written form of informed consent was obtained from all healthy donors (available on request). Blood was collected and methods were carried out in accordance with the relevant guidelines of the ethics committee.

### Sample population

40 healthy individuals were included in the study. Exclusion criteria for the healthy population were: known (chronic and acute) inflammatory conditions such as asthma, human immunodeficiency virus (HIV) or tuberculosis; risk factors associated with metabolic syndrome; smoking; and, if female, being on contraceptive or hormone replacement treatment. This population did not take any anti-inflammatory medication. Whole blood (WB) of the participants was obtained in citrate tubes. WB was used for thromboelastography (TEG) [8, 88] and platelet poor plasma (PPP) was used for confocal and super-resolution analysis.

### Iron, LPS, LTA1 and LTA2

The following concentrations of the various amyloid-inducing substances were used:

- A final exposure iron concentration (FeCl_3_, MW 270.32) of 5 µM was used in all experiments.
- The LPS used was from *E. coli* O111:B4 (Sigma, L2630). A final LPS exposure concentration of 0.4 ng.L^-1^ was used.
- LTA1 was from *Staphylococcus aureus* (Sigma, L2515) and a final LTA1 exposure concentration of 1 ng.L^-1^ was used in all experiments.
- LTA2 was from *Streptococcus pyogenes* (Sigma, L3140) and a final LTA1 exposure concentration of 1 ng.L^-1^ was used in all experiments.

### Confocal microscopy

Platelet poor plasma (PPP) was prepared by centrifuging WB for 15 minutes at 3000 x g, followed by storage at -80°C. On the day of analysis, all -80°C-stored PPPs were brought to room temperature and incubated for one hour with the four candidate amyloidogenic molecules (either LPS, iron, LTA1 or LTA2 (final concentrations given in previous section)) before adding fluorescent markers. This pre-incubation was followed by an incubation of 30 minutes with ThT at a final concentration of 5 µM and Amytracker™ 480 and 680 (0.1 µL into 100 µL PPP). Naïve PPP was incubated only with the 3 markers. Before viewing clots on the confocal microscope, thrombin was added in the ratio 1:2, (5 µL thrombin: 10 µL PPP) and to create extensive fibrin networks, created. Thrombin was provided by the South African National Blood Service, and the thrombin solution was at a concentration of 20 U/ml and made up in a biological buffer containing 0.2% human serum albumin. A coverslip was placed over the prepared clot, and samples were viewed using a Zeiss LSM 780 with ELYRA PS1 confocal microscope with a Plan-Apochromat 63x/1.4 Oil DIC objective. For ThT, the excitation laser used was 488 nm and emission measured at 508 to 570 nm; for Amytracker^TM^ 480 the 405 nm excitation laser was used with emission measured at 478 to 539 nm; and for Amytracker ^TM^ 680 the 561 nm excitation laser was used for excitation with emission measured at 597 to 695 nm. A selection of micrographs of the prepared clots with and without the four molecules was captured. Gain settings were kept the same during all data capture and used for statistical analyses; however, brightness and contrast were slightly adjusted for figure preparation. We also prepared Z-stacks of clots where the candidate amyloidogenic molecules (iron, LPS, LTA1 and LTA2) were added to PPP.

### Thromboelastography

#### Clot property studies

Whole blood (WB) was incubated for 24 hours at room temperature with either iron, LPS, LTA1 or LTA2, or left untreated (naïve sample). Clot property studies using thromboelastography (TEG) were performed as follows: 340µL of naïve or treated WB were placed in a TEG cup and 20µl of 0.2M CaCl_2_ was added. CaCl_2_ is necessary to reverse the effect of the collecting tube’s sodium citrate and consequently initiate coagulation. The samples were then placed in a Thromboelastograph 5000 Hemostasis Analyzer System for analysis. Seven parameters, as shown in Table 1, were studied [88–90].

**Table 1:** TEG clot parameters for whole blood and platelet poor plasma (taken from [88]).

| PARAMETERS | EXPLANATION |
| --- | --- |
| **R value: reaction time measured in minutes** | Time of latency from start of test to initial fibrin formation (amplitude of 2mm); i.e. initiation time |
| **K: kinetics measured in minutes** | Time taken to achieve a certain level of clot strength (amplitude of 20mm); i.e. amplification |
| **A (Alpha): Angle (slope between the traces represented by R and K measured in degrees** | The angle measures the speed at which fibrin build up and cross linking takes place, hence assesses the rate of clot formation; i.e. thrombin burst |
| **MA: Maximal Amplitude measured in mm** | Maximum strength/stiffness of clot. Reflects the ultimate strength of the fibrin clot, i.e. overall stability of the clot |
| **Maximum rate of thrombus generation (MRTG) measured in Dyn.cm<sup>-2</sup>.s<sup>-1</sup>** | The maximum velocity of clot growth observed or maximum rate of thrombus generation using G, where G is the elastic modulus strength of the thrombus in dynes per cm<sup>-2</sup> |
| **Time to maximum rate of thrombus generation (TMRTG) measured in minutes** | The time interval observed before the maximum speed of the clot growth |
| **Total thrombus generation (TTG) measured in Dyn.cm<sup>-2</sup>** | The clot strength: the amount of total resistance (to movement of the cup and pin) generated during clot formation. This is the total area under the velocity curve during clot growth, representing the amount of clot strength generated during clot growth |

### Statistical analysis

TEG results were analysed using the STATSDIRECT (version 2.8.0) software using the paired t-test; notwithstanding its arbitrary nature [91–93], significance was taken as P ≤ 0.05. Confocal techniques are usually used only as qualitative methods. We captured the fluorescent signal of each of the three fluorescent markers as a composite.czi file in the Zeiss ZEN software and then used ImageJ (FIJI) to split the channels. Then we assessed the variance between (black) background and the presence of fluorescent pixels (binary comparison) for each of the three fluorescent markers in clots. For this, we used the histogram function in ImageJ (FIJI) and calculated the coefficient of variation (CV) (as SD/mean) as our metric to quantify and discriminate between clots of healthy naïve PPP and clots with the candidate amyloidogenic molecules. Sample analysis was performed with the Mann-Whitney U test, using the STATSDIRECT (version 2.8.0) software.

### Raw data storage

Raw data are stored on OneDrive that is an open access storage database: (https://1drv.ms/f/s!AgoC0mY3bkKHrx0mNYcZwf2i30w6) and on the first corresponding author’s ResearchGate profile, https://www.researchgate.net/profile/Etheresia_Pretorius

## RESULTS

### Confocal microscopy

Confocal analysis of healthy clotted naïve PPP, in the presence of ThT, Amytracker™ 480 and 680, showed occasional small patches of fluorescence (see Figure 2A to C). However, when any of the four candidate amyloidogenic molecules were pre-incubated with healthy PPP prior to addition of thrombin, the fluorescence in all three channels was greatly enhanced. This suggests increased binding of ThT, as well as of the two Amytracker markers. Amytracker binding, in particular, is a confirmation that amyloidogenesis is promoted by exposure to the four molecules. Amyloidogenesis was most prominent in PPP exposed to the 2 LTAs, suggesting that there are increased β-sheet-rich amyloid areas in the LTA-exposed fibrin(ogen) (Figure 2J to O). Previously, we concluded that LPS binding causes the fibrinogen to polymerise into a form with a greatly increased amount of β-sheet (in the presence of thrombin), reflecting amyloid formation [12]. This results in a strong fluorescence observable (when excited ca 440 nm) in the presence of ThT (see e.g. [11, 35, 36, 49, 94]). Here we confirm that the LPS, iron and the 2 LTAs not only result in ThT binding, plausibly to open hydrophobic areas on fibrin, but that lipoteichoic acids can indeed initiate amyloidogenesis of fibrin(ogen) (as confirmed by the Amytracker™ 480 and 680 binding). The analysis of the micrographs suggest that the Amytracker^TM^ 480 and 680 and the ThT do not bind at identical molecular sites on the fibrin, but that they bind in the same molecular vicinity. We therefore suggest that ThT and Amytracker™ binding do not interfere with each other. Figure 3A to D also show separate and composite z-stack figures of healthy PPP exposed to the 4 different molecules. These results suggest that the Amytrackers^TM^ mainly bind on different parts of the proteins, and that their binding pattern differs between the amyloid formed in the presence of the four different amyloidogenic molecules. Videos of the z-stacks are stored with raw data on OneDrive (see supplementary information). Statistical (coefficient of variation, CV) data are plotted in Figure 4. Data from the three different markers of each of the four amyloidogenic molecules all differed significantly from that of the controls (P ≤ 0.0001). CV analyses were done on over 1750 micrographs.

**Figure 2:**
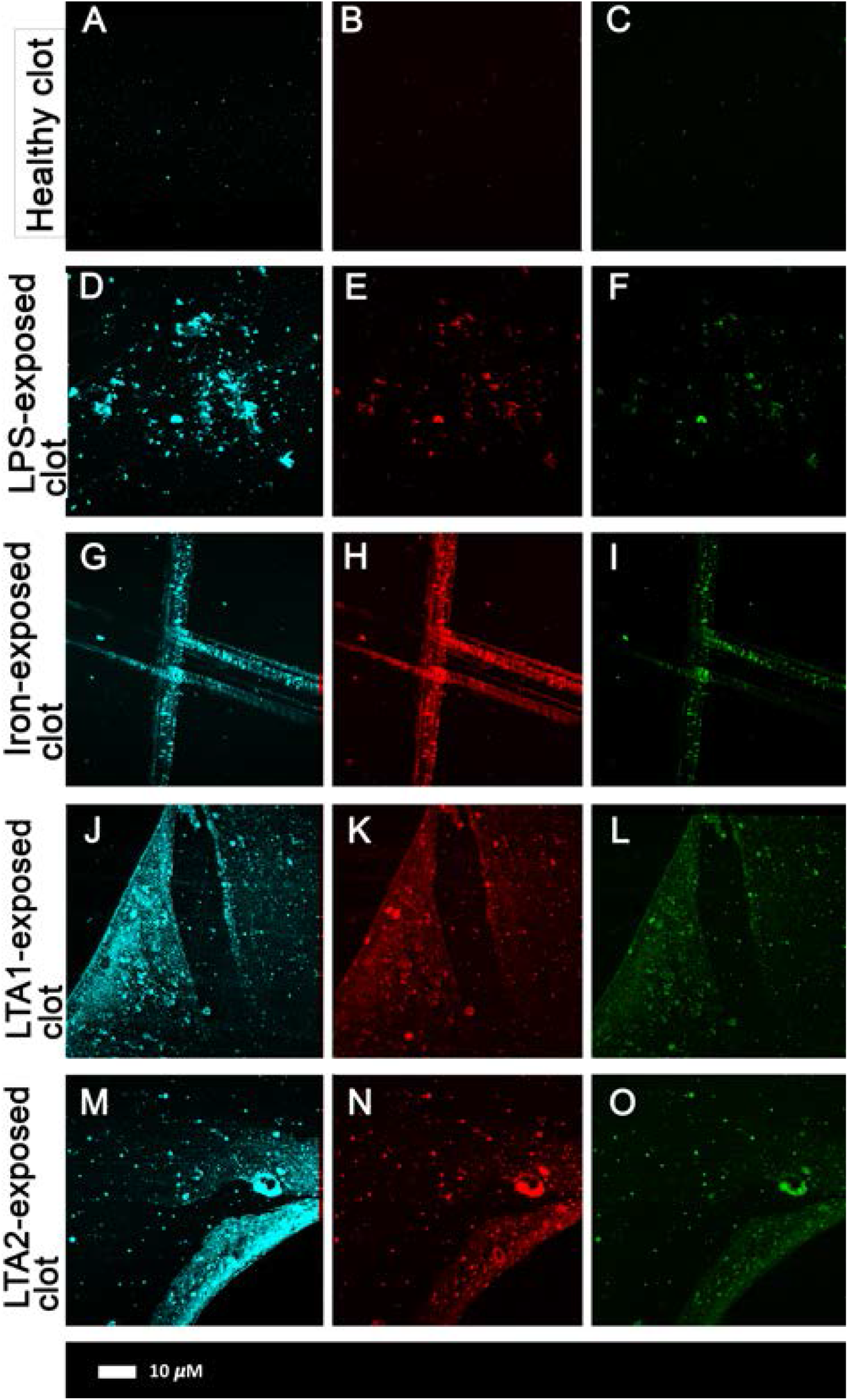
**A)** Representative confocal images of 3 markers (cyan: Amytracker™ 480; red: Amytracker™ 680; green: ThT). The following micrographs are representative of the various exposures: **A to C)** naïve PPP; **D to F)** PPP exposed to LPS; **G to I)** PPP exposed to iron; **J to L)** PPP exposed to LTA1; **M to O)** PPP exposed to LTA2.

**Figure 3:**
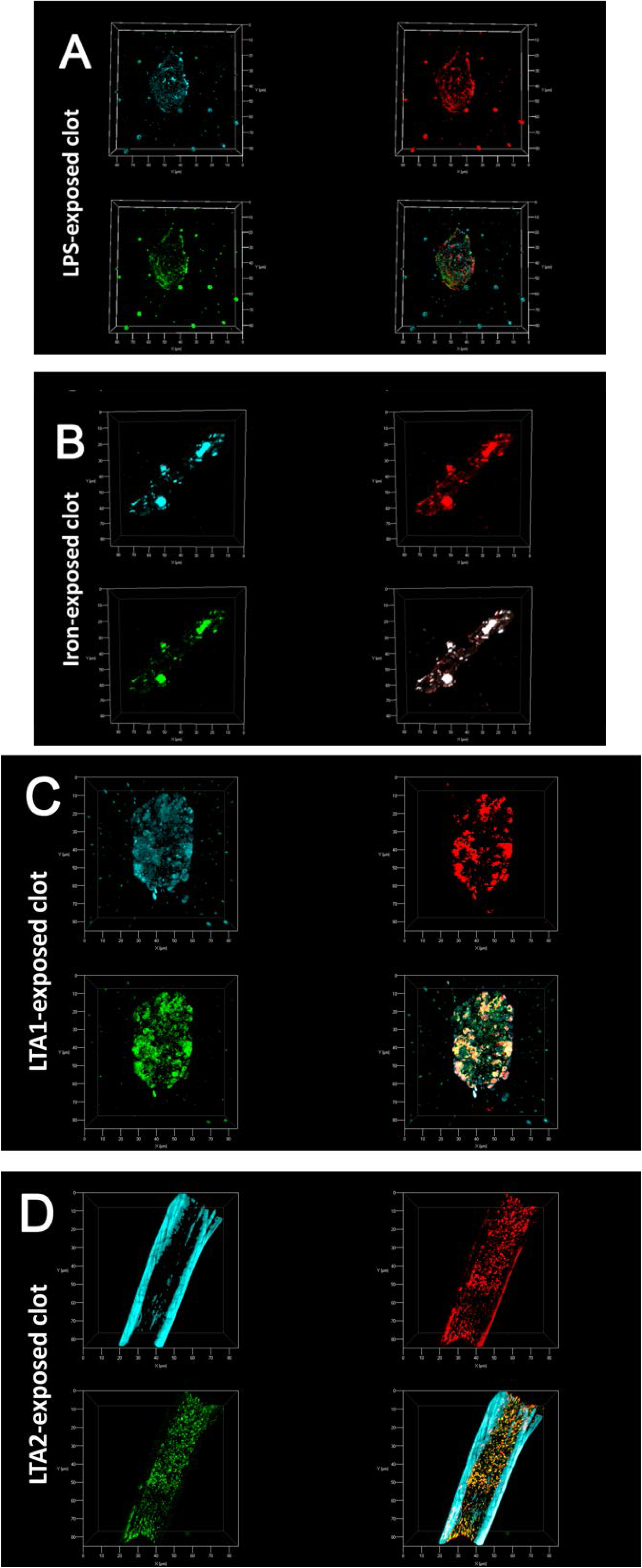
Z-stack projections were created with confocal microscopy and ZEN software by adding four candidate amyloidogenic molecules to naïve plasma. From top left clockwise each figure shows Amytracker™ 480 (cyan); Amytracker™ 680 (red) and ThT (green). Bottom right shows the composite of the 3 markers. Note that in some instances the composite shows white areas; these areas are where all 3 markers overlap. **A)** Clots of PPP exposed to LPS; **B)** Clots of PPP exposed to iron; **C)** Clots of PPP exposed to LTA1; **D)** Clots of PPP exposed to LTA2.

**Figure 4:**
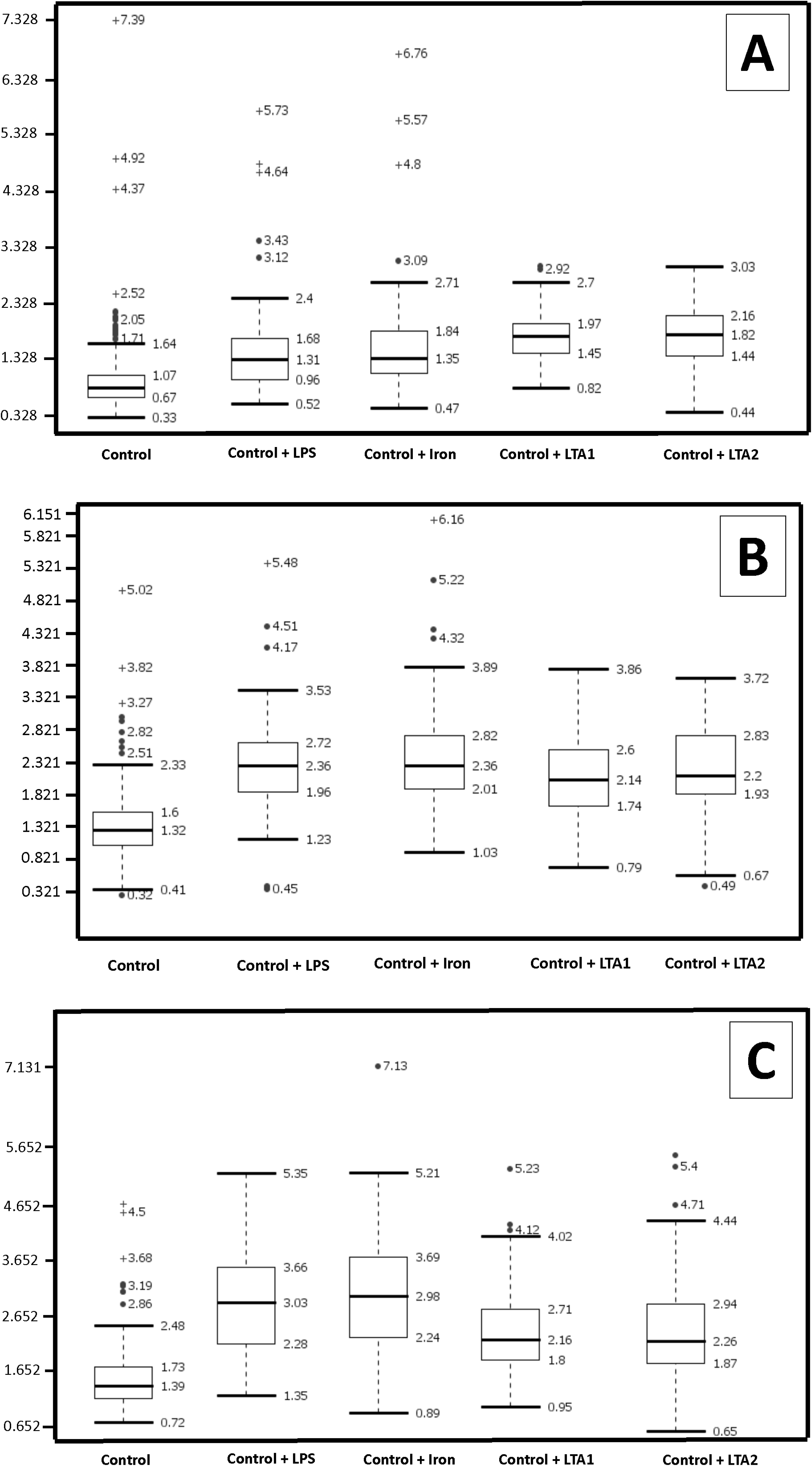
Boxplot of the distribution of the coefficients of variation (CV) for the pixel intensities in the confocal clot images from the three different markers analysed (median coefficients of variation and STDs for each group are reported above the plots). **A)** Amytracker™ 480 **B)** Amytracker™ 680 **C)** ThT. Data from the 3 different markers of each of the 4 molecules all differed significantly from that of the controls (P ≤ 0.0001).

### Thromboelastography

Table 2 shows demographic data of the sample (used for confocal and TEG analysis) and the TEG results of the naïve WB, as well as the results after 24-hour exposure with the 4 different molecules. There is a significantly decreased R-value in the presence of LPS, suggesting that the LPS causes WB to clot faster [88]. This was also previously demonstrated with a lower LPS concentration.

**Table 2:** Demographics and thromboelastography results of naïve blood versus LPS, iron, LTA1 and LTA2. Data in table shows median ± STD for full dataset (n-value of sample size in table header) of TEG parameters for the particular exposure. p-values were calculated by a paired-T-test using only the corresponding naïve sample.

| Healthy Individuals (N=40) |  |  |  |  |  |  |  |  |  |
| --- | --- | --- | --- | --- | --- | --- | --- | --- | --- |
| Gender |  |  |  |  | Age |  |  |  |  |
| 55% F; 45% M |  |  |  |  | 32.5 (±18.8) |  |  |  |  |
| TEG results naïve whole blood after 24 hour exposure to LPS, Iron, LTA1 and LTA2 |  |  |  |  |  |  |  |  |  |
|  | Naïve (n=32) | LPS (n=30) | P-value<br>(Naïve/LPS) | Iron (n=27) | P-value<br>(Naïve/iron) | LTA1<br>(n=15) | P-value<br>(Naïve/LTA 1) | LTA2<br>(n=15) | P-value<br>(Naïve/LTA 2) |
| R | 9.7 ± 1.8 | 8.5 ± 1.6 | 0.02 | 7.1 ± 1.1 | < 0.0001 | 9.2 ± 1.6 | 0.50 | 8.3 ± 1.6 | 0.25 |
| K | 3.1 ± 0.9 | 3.4 ± 1.0 | 0.34 | 2.7 ± 0.8 | 0.003 | 2.7 ± 0.8 | 0.07 | 2.7 ± 0.9 | 0.14 |
| Angle | 50.7 ± 7.8 | 47.6 ± 10.0 | 0.18 | 53.7 ± 7.8 | 0.0004 | 55.9 ± 8.3 | 0.11 | 54.2 ± 8.7 | 0.97 |
| MA | 59.0 ± 7.9 | 57.9 ± 5.8 | 0.38 | 57.9 ± 6.0 | 0.61 | 59.4 ± 7.3 | 0.26 | 60.8 ± 6.2 | 0.007 |
| MRTG | 4.22 ± 1.43 | 4.13 ± 1.26 | 0.29 | 4.39 ± 1.51 | 0.11 | 5.04 ± 2.60 | 0.16 | 5.49 ± 1.82 | 0.11 |
| TMRTG | 13.42 ± 2.70 | 12.54<br>± 2.72 | 0.17 | 10.17 ± 1.67 | < 0.0001 | 12.25 ±<br>2.23 | 0.19 | 11.58 ±<br>2.55 | 0.26 |
| TTG | 729.8 ±<br>247.68 | 689.69± 16<br>3.71 | 0.12 | 702.28 ±<br>170.15 | 0.36 | 729.95 ±<br>246.81 | 0.40 | 779.91 ±<br>205.04 | 0.014 |

Iron results show significant P-values for R, K, angle and TMRTG values, suggesting that the clot forms faster, the clot reaches its (20 mm) set point more slowly, there is more crosslinking of fibrin fibres and there is a decreased time from clot initiation to maximum clot formation. Considering that iron affects more TEG parameters than does LPS, free iron therefore has a more profound effect on clot formation than does LPS at the concentrations used, causing whole blood to be more hypercoagulable [8]. When LTA2 was used, only the MA parameter (maximal amplitude measured in mm) of LTA2 was significantly different from the naïve MA. Overall, the TEG results provide further evidence that the type of amyloid formed differs between LPS, iron, LTA1 and LTA2 when they are used as inducers

## DISCUSSION

In previous work [11, 12], as part of a systems biology approach to understanding the dormant bacterial basis and aetiology of coagulopathies accompanying various inflammatory diseases (e.g. [2, 4, 5, 9, 95–97]), we demonstrated that exceptionally low (and highly substoichiometric) concentrations of LPS could induce the formation of an amyloid form of fibrin. A chief piece of evidence was the extensive fluorescence staining when ThT was added. This was an entirely novel and unexpected finding, although Strickland and colleagues have shown that fibrin(ogen) can interact with known β-amyloid-forming peptides and proteins (e.g. [98–103]).

LPS is a component of Gram-negative bacteria, so an obvious question arose as to whether or not the equivalent molecules in Gram-positive organisms (especially lipoteichoic acid) would have similar effects. In addition, we wished to take the opportunity to assess the utility of various novel conjugated oligothiophene amyloid stains, now commercially available as the ‘Amytracker’™ series, to illuminate amyloids.

In the present paper, we show that LPS, iron, and the 2 LTAs cause amyloid formation of plasma proteins, and in particular of fibrin(ogen) as blood is clotted. We confirmed amyloidogenesis by using small ligands identifying amyloid protein deposits [72]. Specifically, we used Amytracker™ 480 (related to HS163) and 680 (related to HS169) which are fluorescent amyloid ligands, also termed luminescent conjugated oligothiophenes (LCOs) [72, 84]. These two fluorescent markers bind rapidly and with high sensitivity to detect protein amyloid formation in fibrin(ogen). Previously we showed that ThT binds to areas of amyloidogenic fibrin(ogen) that are created by LPS exposure during clotting. Here we confirm that observation using LPS, and show further that iron and the 2 LTAs all cause changes in fibrin(ogen) conformation to an amyloid(ogenic) nature, with the role of iron being recognised in assisting the regrowth of dormant bacteria that can then shed the inflammagenic cell wall materials [11].

It is known that the Amytracker™ dyes are spectrally richer, can discriminate different forms of amyloid, and that their staining properties clearly differ from those of ThT [68, 70, 77–79]. Comparison (e.g. Figure 2) of the staining with the three dyes (ThT and Amytracker™ 480 and 680) in the presence of the four candidate amyloidogenic molecules showed that this is also true for the amyloid form(s) of fibrin induced by the different agents. These clearly different staining patterns occur for each dye, with those of the two Amytracker^TM^ dyes showing greater staining and being more similar (but not identical) to each other. Because the stoichiometry is of the order of 10^−8^ LPS/LTA:fibrinogen, we have been unable to determine the bindings sites, though clear the fact that they differ is the underlying cause of the different morphologies observed. As a referee pointed out, it is possible to view the binding of LPS/LTA by fibrinogen as a host-protective mechanism; however, the balance between the inflammatory potential of LPS/LTA and the amyloid forms of fibrin is, as yet, unknown.

This becomes even clearer when we observe the staining in the z-stack projections (Figure 3 and supplementary video information of the clots shown in Figure 3), with the Amytracker™ 480 seeming to favour the larger fibres characteristic of LTA2. These differences were also observed in the TEG traces: while iron changed four of the TEG variables significantly (R, K, angle and TMRTG), LPS showed a significantly increased Rvalue, LTA1 was without effect, while LTA2 affected only the MA (and that marginally).

We conclude that lipoteichoic acids are even more potent and effective than is LPS in binding to fibrinogen and in affecting the manner in which it self-organises during blood clotting. Such findings have profound significance for our understanding of the aetiology of anomalous blood clots, and may have value in diagnosis, prognosis and treatment of chronic, inflammatory diseases.

## ACKNOWLEDGMENTS

We thank the Biotechnology and Biological Sciences Research Council (grant BB/L025752/1), as well as the National Research Foundation (NRF) and Medical Research Council (MRC) of South Africa for supporting this collaboration. We also thank the SANBS for providing the human thrombin without cost.

## DECLARATION OF AUTHORSHIP

EP: co-project leader, confocal and statistical analysis; MJP: figure analysis and rendering, technical preparation of all samples, TEG analysis; SRB, NBN, LH: technical assistance; DBK: co-project leader and originator of suggestion of using LTAs.

## SUPPLEMENTARY MATERIAL

Z-stack videos were created with Zeiss ZEN software and stored as supplementary material on OneDrive that is an open access storage database: (https://1drv.ms/f/s!AgoC0mY3bkKHrx0mNYcZwf2i30w6) as well as within the publisher link to the paper. When the video is played of each healthy clot where the LPS, iron, LTA1 and LTA2 were added, different binding areas of the 3 markers are clearly visible.

## REFERENCES

[1] Kell, D.B., Potgieter, M. & Pretorius, E. 2015 Individuality, phenotypic differentiation, dormancy and ‘persistence’ in culturable bacterial systems: commonalities shared by environmental, laboratory, and clinical microbiology. F1000Research 4, 179. (doi:10.12688/f1000research.6709.1).

[2] Kell, D.B. & Pretorius, E. 2015 On the translocation of bacteria and their lipopolysaccharides between blood and peripheral locations in chronic, inflammatory diseases: the central roles of LPS and LPS-induced cell death Integr Biol 7, 1339–1377. (doi:10.1039/C5IB00158G).

[3] Potgieter, M., Bester, J., Kell, D.B. & Pretorius, E. 2015 The dormant blood microbiome in chronic, inflammatory diseases. FEMS Microbiol Rev 39, 567–591. (doi:http://dx.doi.org/10.1093/femsre/fuv013).

[4] Pretorius, E., Akeredolu, O.-O., Soma, P. & Kell, D.B. 2017 Major involvement of bacterial components in rheumatoid arthritis and its accompanying oxidative stress, systemic inflammation and hypercoagulability. Exp Biol Med 242, 355–373.

[5] Pretorius, E., Bester, J. & Kell, D.B. 2016 A bacterial component to Alzheimer-type dementia seen via a systems biology approach that links iron dysregulation and inflammagen shedding to disease J Alzheimers Dis 53, 1237–1256.

[6] Kell, D.B. & Pretorius, E. 2017 To what extent are the terminal stages of sepsis, septic shock, SIRS, and multiple organ dysfunction syndrome actually driven by a toxic prion/amyloid form of fibrin?. Seminars in Thrombosis and Hemostasis In Press.

[7] Kell, D.B. & Pretorius, E. 2014 Serum ferritin is an important disease marker, and is mainly a leakage product from damaged cells. Metallomics 6, 748–773. (doi:10.1039/C3MT00347G).

[8] Kell, D.B. & Pretorius, E. 2015 The simultaneous occurrence of both hypercoagulability and hypofibrinolysis in blood and serum during systemic inflammation, and the roles of iron and fibrin(ogen). Integr Biol 7, 24–52. (doi:10.1039/c4ib00173g).

[9] Pretorius, E. & Kell, D.B. 2014 Diagnostic morphology: biophysical indicators for iron-driven inflammatory diseases. Integrative Biol 6, 486–510.

[10] Pretorius, E., Vermeulen, N., Bester, J., Lipinski, B. & Kell, D.B. 2013 A novel method for assessing the role of iron and its functional chelation in fibrin fibril formation: the use of scanning electron microscopy. Toxicol Mech Methods 23, 352–359. (doi:10.3109/15376516.2012.762082).

[11] Kell, D.B. & Pretorius, E. 2017 Proteins behaving badly. Substoichiometric molecular control and amplification of the initiation and nature of amyloid fibril formation: lessons from and for blood clotting. Progr Biophys Mol Biol 123, 16–41. (doi:http://dx.doi.org/10.1016/j.pbiomolbio.2016.08.006).

[12] Pretorius, E., Mbotwe, S., Bester, J., Robinson, C.J. & Kell, D.B. 2016 Acute induction of anomalous and amyloidogenic blood clotting by molecular amplification of highly substoichiometric levels of bacterial lipopolysaccharide. J R Soc Interface 123, 20160539. (doi:http://dx.doi.org/10.1098/rsif.2016.0539).

[13] Schwandner, R., Dziarski, R., Wesche, H., Rothe, M. & Kirschning, C.J. 1999 Peptidoglycan-and lipoteichoic acid-induced cell activation is mediated by toll-like receptor 2. J Biol Chem 274, 17406–17409.

[14] Morath, S., Geyer, A. & Hartung, T. 2001 Structure-function relationship of cytokine induction by lipoteichoic acid from *Staphylococcus aureus*. J Exp Med 193, 393–397.

[15] Poltorak, A., He, X.L., Smirnova, I., Liu, M.Y., Van Huffel, C., Du, X., Birdwell, D., Alejos, E., Silva, M., Galanos, C., et al. 1998 Defective LPS signaling in C3H/HeJ and C57BL/10ScCr mice: Mutations in Tlr4 gene. Science 282, 2085–2088. (doi:DOI 10.1126/science.282.5396.2085).

[16] Hoshino, K., Takeuchi, O., Kawai, T., Sanjo, H., Ogawa, T., Takeda, Y., Takeda, K. & Akira, S. 1999 Toll-like receptor 4 (TLR4)-deficient mice are hyporesponsive to lipopolysaccharide: Evidence for TLR4 as the Lps gene product. J Immunol 162, 3749–3752.

[17] Lien, E., Means, T.K., Heine, H., Yoshimura, A., Kusumoto, S., Fukase, K., Fenton, M.J., Oikawa, M., Qureshi, N., Monks, B., et al. 2000 Toll-like receptor 4 imparts ligand-specific recognition of bacterial lipopolysaccharide. J Clin Invest 105, 497–504. (doi:10.1172/JCI8541).

[18] Underhill, D.M., Ozinsky, A., Hajjar, A.M., Stevens, A., Wilson, C.B., Bassetti, M. & Aderem, A. 1999 The Toll-like receptor 2 is recruited to macrophage phagosomes and discriminates between pathogens. Nature 401, 811–815. (doi:10.1038/44605).

[19] Ishii, K.J. & Akira, S. 2004 Toll-like Receptors and Sepsis. Curr Infect Dis Rep 6, 361366.

[20] Zähringer, U., Lindner, B., Inamura, S., Heine, H. & Alexander, C. 2008 TLR2 - promiscuous or specific? A critical re-evaluation of a receptor expressing apparent broad specificity. Immunobiology 213, 205–224. (doi:10.1016/j.imbio.2008.02.005).

[21] Kawai, T. & Akira, S. 2011 Toll-like receptors and their crosstalk with other innate receptors in infection and immunity. Immunity 34, 637–650. (doi:DOI 10.1016/j.immuni.2011.05.006).

[22] Kumar, H., Kawai, T. & Akira, S. 2011 Pathogen recognition by the innate immune system. Int Rev Immunol 30, 16–34. (doi:10.3109/08830185.2010.529976).

[23] Oliveira-Nascimento, L., Massari, P. & Wetzler, L.M. 2012 The Role of TLR2 in Infection and Immunity. Front Immunol 3, 79. (doi:10.3389/fimmu.2012.00079).

[24] Kumar, S., Ingle, H., Prasad, D.V. & Kumar, H. 2013 Recognition of bacterial infection by innate immune sensors. Crit Rev Microbiol 39, 229–246. (doi:10.3109/1040841X.2012.706249).

[25] Alexander, S.P.A., Benson, H.E., Faccenda, E., Pawson, A.J., Sharman, J.L., Spedding, M., Peters, J.A., Harmar, A.J. & CGTP Collaborators. 2013 The Concise Guide to PHARMACOLOGY 2013/14: catalytic receptors. Br J Pharmacol 170, 1676–1705. (doi:10.1111/bph.12449).

[26] Liu, Y., Yin, H., Zhao, M. & Lu, Q. 2014 TLR2 and TLR4 in autoimmune diseases: a comprehensive review. Clin Rev Allergy Immunol 47, 136–147. (doi:10.1007/s12016-013-8402-y).

[27] Mukherjee, S., Karmakar, S. & Babu, S.P. 2016 TLR2 and TLR4 mediated host immune responses in major infectious diseases: a review. Braz J Infect Dis 20, 193–204. (doi: 10.1016/j.bjid.2015.10.011).

[28] Jiménez-Dalmaroni, M.J., Gerswhin, M.E. & Adamopoulos, I.E. 2016 The critical role of toll-like receptors-From microbial recognition to autoimmunity: A comprehensive review. Autoimmun Rev 15, 1–8. (doi:10.1016/j.autrev.2015.08.009).

[29] Morath, S., von Aulock, S. & Hartung, T. 2005 Structure/function relationships of lipoteichoic acids. J Endotoxin Res 11, 348–356. (doi:10.1179/096805105X67328).

[30] Ninkovic, J., Anand, V., Dutta, R., Zhang, L., Saluja, A., Meng, J., Koodie, L., Banerjee, S. & Roy, S. 2016 Differential effects of gram-positive and gram-negative bacterial products on morphine induced inhibition of phagocytosis. Sci Rep 6, 21094. (doi:10.1038/srep21094).

[31] Hoogerwerf, J.J., de Vos, A.F., Levi, M., Bresser, P., van der Zee, J.S., Draing, C., von Aulock, S. & van der Poll, T. 2009 Activation of coagulation and inhibition of fibrinolysis in the human lung on bronchial instillation of lipoteichoic acid and lipopolysaccharide. Crit Care Med 37, 619–625. (doi:10.1097/CCM.0b013e31819584f9).

[32] Barbero-Becerra, V.J., Gutiérrez-Ruiz, M.C., Maldonado-Bernal, C., Téllez-Avila, F.I., Alfaro-Lara, R. & Vargas-Vorácková, F. 2011 Vigorous, but differential mononuclear cell response of cirrhotic patients to bacterial ligands. World J Gastroenterol 17, 1317–1325. (doi:10.3748/wjg.v17.i10.1317).

[33] Cinar, M.U., Islam, M.A., Pröll, M., Kocamis, H., Tholen, E., Tesfaye, D., Looft, C., Schellander, K. & Uddin, M.J. 2013 Evaluation of suitable reference genes for gene expression studies in porcine PBMCs in response to LPS and LTA. BMC Res Notes 6, 56. (doi: 10.1186/1756-0500-6-56).

[34] Biancalana, M., Makabe, K., Koide, A. & Koide, S. 2009 Molecular mechanism of thioflavin-T binding to the surface of beta-rich peptide self-assemblies. J Mol Biol 385, 1052–1063. (doi:10.1016/j.jmb.2008.11.006).

[35] Biancalana, M. & Koide, S. 2010 Molecular mechanism of Thioflavin-T binding to amyloid fibrils. Biochim Biophys Acta 1804, 1405–1412. (doi:10.1016/j.bbapap.2010.04.001).

[36] Groenning, M. 2010 Binding mode of Thioflavin T and other molecular probes in the context of amyloid fibrils-current status. J Chem Biol 3, 1–18. (doi:10.1007/s12154-009-0027-5).

[37] Amdursky, N., Erez, Y. & Huppert, D. 2012 Molecular rotors: what lies behind the high sensitivity of the thioflavin-T fluorescent marker. Acc Chem Res 45, 1548–1557. (doi:10.1021/ar300053p).

[38] Khurana, R., Coleman, C., Ionescu-Zanetti, C., Carter, S.A., Krishna, V., Grover, R.K., Roy, R. & Singh, S. 2005 Mechanism of thioflavin T binding to amyloid fibrils. J Struct Biol 151, 229–238. (doi:10.1016/j.jsb.2005.06.006).

[39] Krebs, M.R.H., Bromley, E.H. & Donald, A.M. 2005 The binding of thioflavin-T to amyloid fibrils: localisation and implications. J Struct Biol 149, 30–37. (doi:10.1016/j.jsb.2004.08.002).

[40] LeVine, H., 3rd. 1999 Quantification of beta-sheet amyloid fibril structures with thioflavin T. Methods Enzymol 309, 274–284.

[41] Ivancic, V., Ekanayake, O. & Lazo, N. 2016 Binding Modes of Thioflavin T on the Surface of Amyloid Fibrils by NMR. ChemPhysChem. (doi:10.1002/cphc.201600246).

[42] Lindberg, D.J., Wranne, M.S., Gilbert Gatty, M., Westerlund, F. & Esbjörner, E.K. 2015 Steady-state and time-resolved Thioflavin-T fluorescence can report on morphological differences in amyloid fibrils formed by Abeta(1-40) and Abeta(1-42). Biochem Biophys Res Commun 458, 418–423. (doi:10.1016/j.bbrc.2015.01.132).

[43] Murugan, N.A., Olsen, J.M.H., Kongsted, J., Rinkevicius, Z., Aidas, K. & Ågren, H. 2013 Amyloid Fibril-Induced Structural and Spectral Modifications in the Thioflavin-T Optical Probe. J Phys Chem Lett 4, 70–77. (doi:10.1021/jz3018557).

[44] Picken, M.M. & Herrera, G.A. 2012 Thioflavin T Stain: An Easier and More Sensitive Method for Amyloid Detection. Curr Clin Pathol, 187–189.

[45] Rybicka, A., Longhi, G., Castiglioni, E., Abbate, S., Dzwolak, W., Babenko, V. & Pecul, M. 2016 Thioflavin T: Electronic Circular Dichroism and Circularly Polarized Luminescence Induced by Amyloid Fibrils. Chemphyschem. (doi:10.1002/cphc.201600235).

[46] Sabate, R., Rodriguez-Santiago, L., Sodupe, M., Saupe, S.J. & Ventura, S. 2013 Thioflavin-T excimer formation upon interaction with amyloid fibers. Chem Commun (Camb) 49, 5745–5747. (doi:10.1039/c3cc42040j).

[47] Sulatskaya, A.I., Kuznetsova, I.M. & Turoverov, K.K. 2011 Interaction of thioflavin T with amyloid fibrils: stoichiometry and affinity of dye binding, absorption spectra of bound dye. J Phys Chem B 115, 11519–11524. (doi:10.1021/jp207118x).

[48] Sulatskaya, A.I., Kuznetsova, I.M. & Turoverov, K.K. 2012 Interaction of thioflavin T with amyloid fibrils: fluorescence quantum yield of bound dye. J Phys Chem B 116, 2538–2544. (doi:10.1021/jp2083055).

[49] Younan, N.D. & Viles, J.H. 2015 A Comparison of Three Fluorophores for the Detection of Amyloid Fibers and Prefibrillar Oligomeric Assemblies. ThT (Thioflavin T); ANS (1-Anilinonaphthalene-8-sulfonic Acid); and bisANS (4,4’-Dianilino-1,1’-binaphthyl-5,5’-disulfonic Acid). Biochemistry 54, 4297–4306. (doi:10.1021/acs.biochem.5b00309).

[50] Zhang, X. & Ran, C. 2013 Dual functional small molecule probes as fluorophore and ligand for misfolding proteins. Curr Org Chem 17, 6. (doi:10.2174/1385272811317060004).

[51] Maezawa, I., Hong, H.S., Liu, R., Wu, C.Y., Cheng, R.H., Kung, M.P., Kung, H.F., Lam, K.S., Oddo, S., Laferla, F.M., et al. 2008 Congo red and thioflavin-T analogs detect Abeta oligomers. J Neurochem 104, 457–468. (doi:10.1111/j.1471-4159.2007.04972.x).

[52] Chang, W.M., Dakanali, M., Capule, C.C., Sigurdson, C.J., Yang, J. & Theodorakis, E.A. 2011 ANCA: A Family of Fluorescent Probes that Bind and Stain Amyloid Plaques in Human Tissue. ACS Chem Neurosci 2, 249–255. (doi:10.1021/cn200018v).

[53] Giorgadze, T.A., Shiina, N., Baloch, Z.W., Tomaszewski, J.E. & Gupta, P.K. 2004 Improved detection of amyloid in fat pad aspiration: an evaluation of Congo red stain by fluorescent microscopy. Diagn Cytopathol 31, 300–306. (doi:10.1002/dc.20131).

[54] Nilsson, K.P.R., Hammarström, P., Ahlgren, F., Herland, A., Schnell, E.A., Lindgren, M., Westermark, G.T. & Inganäs, O. 2006 Conjugated polyelectrolytes–conformation-sensitive optical probes for staining and characterization of amyloid deposits. ChemBioChem 7, 1096-1104. (doi:10.1002/cbic.200500550).

[55] Rajamohamedsait, H.B. & Sigurdsson, E.M. 2012 Histological staining of amyloid and pre-amyloid peptides and proteins in mouse tissue. Methods Mol Biol 849, 411–424. (doi:10.1007/978-1-61779-551 -0_28).

[56] Volkova, K.D., Kovalska, V.B., Balanda, A.O., Vermeij, R.J., Subramaniam, V., Slominskii, Y.L. & Yarmoluk, S.M. 2007 Cyanine dye-protein interactions: looking for fluorescent probes for amyloid structures. J Biochem Biophys Methods 70, 727–733. (doi:10.1016/j.jbbm.2007.03.008).

[57] Mishra, R., Sjolander, D. & Hammarström, P. 2011 Spectroscopic characterization of diverse amyloid fibrils *in vitro* by the fluorescent dye Nile red. Mol Biosyst 7, 1232–1240. (doi:10.1039/c0mb00236d).

[58] Shen, D., Coleman, J., Chan, E., Nicholson, T.P., Dai, L., Sheppard, P.W. & Patton, W.F. 2011 Novel cell-and tissue-based assays for detecting misfolded and aggregated protein accumulation within aggresomes and inclusion bodies. Cell Biochem Biophys 60, 173–185. (doi:10.1007/s12013-010-9138-4).

[59] Easterhoff, D., DiMaio, J.T.M., Liyanage, W., Lo, C.W., Bae, W., Doran, T.M., Smrcka, A., Nilsson, B.L. & Dewhurst, S. 2013 Fluorescence detection of cationic amyloid fibrils in human semen. Bioorg Med Chem Lett 23, 5199–5202. (doi:10.1016/j.bmcl.2013.06.097).

[60] Kovalska, V.B., Losytskyy, M.Y., Tolmachev, O.I., Slominskii, Y.L., Segers-Nolten, G.M., Subramaniam, V. & Yarmoluk, S.M. 2012 Tri-and pentamethine cyanine dyes for fluorescent detection of alpha-synuclein oligomeric aggregates. J Fluoresc 22, 1441–1448. (doi:10.1007/s10895-012-1081-x).

[61] Ono, M., Watanabe, H., Kimura, H. & Saji, H. 2012 BODIPY-based molecular probe for imaging of cerebral beta-amyloid plaques. ACS Chem Neurosci 3, 319–324. (doi: 10.1021/cn3000058).

[62] Rajasekhar, K., Narayanaswamy, N., Murugan, N.A., Kuang, G., Agren, H. & Govindaraju, T. 2016 A High Affinity Red Fluorescence and Colorimetric Probe for Amyloid beta Aggregates. Sci Rep 6, 23668. (doi:10.1038/srep23668).

[63] Watanabe, H., Ono, M., Matsumura, K., Yoshimura, M., Kimura, H. & Saji, H. 2013 Molecular imaging of beta-amyloid plaques with near-infrared boron dipyrromethane (BODIPY)-based fluorescent probes. Mol Imaging 12, 338–347.

[64] Yuan, L., Lin, W., Zheng, K., He, L. & Huang, W. 2013 Far-red to near infrared analyte-responsive fluorescent probes based on organic fluorophore platforms for fluorescence imaging. Chem Soc Rev 42, 622–661. (doi:10.1039/c2cs35313j).

[65] Guo, Z., Park, S., Yoon, J. & Shin, I. 2014 Recent progress in the development of near-infrared fluorescent probes for bioimaging applications. Chem Soc Rev 43, 16–29. (doi:10.1039/c3cs60271k).

[66] Staderini, M., Martin, M.A., Bolognesi, M.L. & Menéndez, J.C. 2015 Imaging of beta-amyloid plaques by near infrared fluorescent tracers: a new frontier for chemical neuroscience. Chem Soc Rev 44, 1807–1819. (doi:10.1039/c4cs00337c).

[67] Zhang, X., Tian, Y., Zhang, C., Tian, X., Ross, A.W., Moir, R.D., Sun, H., Tanzi, R.E., Moore, A. & Ran, C. 2015 Near-infrared fluorescence molecular imaging of amyloid beta species and monitoring therapy in animal models of Alzheimer’s disease. Proc Natl Acad Sci U S A 112, 9734–9739. (doi:10.1073/pnas.1505420112).

[68] Åslund, A., Sigurdson, C.J., Klingstedt, T., Grathwohl, S., Bolmont, T., Dickstein, D.L., Glimsdal, E., Prokop, S., Lindgren, M., Konradsson, P., et al. 2009 Novel pentameric thiophene derivatives for in vitro and in vivo optical imaging of a plethora of protein aggregates in cerebral amyloidoses. ACS Chem Biol 4, 673–684. (doi:10.1021/cb900112v).

[69] Berg, I., Nilsson, K.P.R., Thor, S. & Hammarström, P. 2010 Efficient imaging of amyloid deposits in *Drosophila* models of human amyloidoses. Nat Protoc 5, 935–944. (doi:10.1038/nprot.2010.41).

[70] Klingstedt, T., Åslund, A., Simon, R.A., Johansson, L.B.G., Mason, J.J., Nyström, S., Hammarström, P. & Nilsson, K.P.R. 2011 Synthesis of a library of oligothiophenes and their utilization as fluorescent ligands for spectral assignment of protein aggregates. Org Biomol Chem 9, 8356–8370. (doi:10.1039/c1ob05637a).

[71] Klingstedt, T. & Nilsson, K.P. 2012 Luminescent conjugated poly-and oligo-thiophenes: optical ligands for spectral assignment of a plethora of protein aggregates. Biochem Soc Trans 40, 704–710. (doi:10.1042/bst20120009).

[72] Klingstedt, T., Blechschmidt, C., Nogalska, A., Prokop, S., Haggqvist, B., Danielsson, O., Engel, W.K., Askanas, V., Heppner, F.L. & Nilsson, K.P.R. 2013 Luminescent conjugated oligothiophenes for sensitive fluorescent assignment of protein inclusion bodies. Chembiochem 14, 607–616. (doi:10.1002/cbic.201200731).

[73] Klingstedt, T., Shirani, H., Åslund, K.O.A., Cairns, N.J., Sigurdson, C.J., Goedert, M. & Nilsson, K.P.R. 2013 The structural basis for optimal performance of oligothiophene-based fluorescent amyloid ligands: conformational flexibility is essential for spectral assignment of a diversity of protein aggregates. Chemistry 19, 10179–10192. (doi: 10.1002/chem.201301463).

[74] Magnusson, K., Simon, R., Sjölander, D., Sigurdson, C.J., Hammarström, P. & Nilsson, K.P.R. 2014 Multimodal fluorescence microscopy of prion strain specific PrP deposits stained by thiophene-based amyloid ligands. Prion 8, 319–329. (doi:10.4161/pri.29239).

[75] Nilsson, K.P., Lindgren, M. & Hammarström, P. 2012 A pentameric luminescent-conjugated oligothiophene for optical imaging of in vitro-formed amyloid fibrils and protein aggregates in tissue sections. Methods Mol Biol 849, 425–434. (doi:10.1007/978-1-61779-551-0_29).

[76] Nyström, S., Psonka-Antonczyk, K.M., Ellingsen, P.G., Johansson, L.B., Reitan, N., Handrick, S., Prokop, S., Heppner, F.L., Wegenast-Braun, B.M., Jucker, M., et al. 2013 Evidence for age-dependent in vivo conformational rearrangement within Abeta amyloid deposits. ACS Chem Biol 8, 1128–1133. (doi:10.1021/cb4000376).

[77] Psonka-Antonczyk, K.M., Duboisset, J., Stokke, B.T., Zako, T., Kobayashi, T., Maeda, M., Nyström, S., Mason, J., Hammarström, P., Nilsson, K.P.R., et al. 2012 Nanoscopic and photonic ultrastructural characterization of two distinct insulin amyloid states. Int J Mol Sci 13, 1461–1480. (doi:10.3390/ijms13021461).

[78] Simon, R.A., Shirani, H., Åslund, K.O.A., Bäck, M., Haroutunian, V., Gandy, S. & Nilsson, K.P.R. 2014 Pentameric thiophene-based ligands that spectrally discriminate amyloid-beta and tau aggregates display distinct solvatochromism and viscosity-induced spectral shifts. Chemistry 20, 12537–12543. (doi:10.1002/chem.201402890).

[79] Sigurdson, C.J., Nilsson, K.P., Hornemann, S., Manco, G., Polymenidou, M., Schwarz, P., Leclerc, M., Hammarström, P., Wüthrich, K. & Aguzzi, A. 2007 Prion strain discrimination using luminescent conjugated polymers. Nat Methods 4, 1023–1030. (doi:10.1038/nmeth1131).

[80] Herrmann, U.S., Schütz, A.K., Shirani, H., Huang, D., Saban, D., Nuvolone, M., Li, B., Ballmer, B., Åslund, A.K.O., Mason, J.J., et al. 2015 Structure-based drug design identifies polythiophenes as antiprion compounds. Sci Transl Med 7, 299ra123. (doi:10.1126/scitranslmed.aab1923).

[81] Margalith, I., Suter, C., Ballmer, B., Schwarz, P., Tiberi, C., Sonati, T., Falsig, J., Nyström, S., Hammarström, P., Åslund, A., et al. 2012 Polythiophenes inhibit prion propagation by stabilizing prion protein (PrP) aggregates. J Biol Chem 287, 18872–18887. (doi:10.1074/jbc.M 112.355958).

[82] Lucanic, M., Plummer, W.T., Chen, E., Harke, J., Foulger, A.C., Onken, B., Coleman-Hulbert, A.L., Dumas, K.J., Guo, S., Johnson, E., et al. 2017 Impact of genetic background and experimental reproducibility on identifying chemical compounds with robust longevity effects. Nat Commun 8, 14256. (doi:10.1038/ncomms14256).

[83] Weininger, D. 1988 SMILES, a chemical language and information system .1. Introduction to methodology and encoding rules. J. Chem. Inf. Comput. Sci. 28, 31–36.

[84] Shirani, H., Linares, M., Sigurdson, C.J., Lindgren, M., Norman, P. & Nilsson, K.P.R. 2015 A Palette of Fluorescent Thiophene-Based Ligands for the Identification of Protein Aggregates. Chemistry 21, 15133–15137. (doi:10.1002/chem.201502999).

[85] Kell, D.B. 2009 Iron behaving badly: inappropriate iron chelation as a major contributor to the aetiology of vascular and other progressive inflammatory and degenerative diseases. BMC Med Genom 2, 2.

[86] Kell, D.B. 2010 Towards a unifying, systems biology understanding of large-scale cellular death and destruction caused by poorly liganded iron: Parkinson’s, Huntington’s, Alzheimer’s, prions, bactericides, chemical toxicology and others as examples. Arch Toxicol 577, 825–889 (doi:10.1007/s00204-010-0577-x).

[87] Pretorius, E., Page, M.J., Hendricks, L., Nkosi, B.N., Benson, S.R. & Kell, D.B. 2017 Both lipopolysaccharide and lipoteichoic acids potently induce anomalous fibrin amyloid formation: assessment with novel Amytracker™ stains. bioRxiv preprint. bioRxiv, 143867.

[88] Pretorius, E., Swanepoel, A.C., DeVilliers, S. & Bester, J. 2017 Blood clot parameters: Thromboelastography and scanning electron microscopy in research and clinical practice. Thromb Res 154, 59–63. (doi:10.1016/j.thromres.2017.04.005).

[89] Bester, J., Soma, P., Kell, D.B. & Pretorius, E. 2015 Viscoelastic and ultrastructural characteristics of whole blood and plasma in Alzheimer-type dementia, and the possible role of bacterial lipopolysaccharides (LPS). Oncotarget 6, 35284–35303.

[90] de Villiers, S., Swanepoel, A., Bester, J. & Pretorius, E. 2015 Novel Diagnostic and Monitoring Tools in Stroke: an Individualized Patient-Centered Precision Medicine Approach. J Atheroscler Thromb 23, 493–504. (doi:10.5551/jat.32748).

[91] Broadhurst, D. & Kell, D.B. 2006 Statistical strategies for avoiding false discoveries in metabolomics and related experiments. Metabolomics 2, 171–196. (doi:10.1007/s11306-006-0037-z).

[92] Greenland, S., Senn, S.J., Rothman, K.J., Carlin, J.B., Poole, C., Goodman, S.N. & Altman, D.G. 2016 Statistical tests, P values, confidence intervals, and power: a guide to misinterpretations. Eur J Epidemiol 31, 337–350. (doi:10.1007/s10654-016-0149-3).

[93] Altman, N. & Krzywinski, M. 2017 Interpreting P values. Nat Meth 14, 213–214.

[94] Freire, S., de Araujo, M.H., Al-Soufi, W. & Novo, M. 2014 Photophysical study of Thioflavin T as fluorescence marker of amyloid fibrils. Dyes and Pigments 110, 97–105. (doi:10.1016/j.dyepig.2014.05.004).

[95] Kell, D.B. & Pretorius, E. 2016 To what extent are the terminal stages of sepsis, septic shock, SIRS, and multiple organ dysfunction syndrome actually driven by a toxic prion/amyloid form of fibrin? bioRxiv preprint. bioRxiv, 057851. (doi:http://dx.doi.org/10.1101/057851).

[96] Kell, D.B. & Kenny, L.C. 2016 A dormant microbial component in the development of pre-eclampsia. Front Med Obs Gynecol 3, 60. (doi:10.3389/fmed.2016.00060).

[97] Pretorius, E., Bester, J., Vermeulen, N., Alummoottil, S., Soma, P., Buys, A.V. & Kell, D. B. 2015 Poorly controlled type 2 diabetes is accompanied by significant morphological and ultrastructural changes in both erythrocytes and in thrombin-generated fibrin: implications for diagnostics. Cardiovasc Diabetol 13, 30.

[98] Ahn, H.J., Zamolodchikov, D., Cortes-Canteli, M., Norris, E.H., Glickman, J.F. & Strickland, S. 2010 Alzheimer’s disease peptide beta-amyloid interacts with fibrinogen and induces its oligomerization. Proc Natl Acad Sci 107, 21812–21817. (doi:10.1073/pnas.1010373107).

[99] Ahn, H.J., Glickman, J.F., Poon, K.L., Zamolodchikov, D., Jno-Charles, O.C., Norris, E. H. & Strickland, S. 2014 A novel Abeta-fibrinogen interaction inhibitor rescues altered thrombosis and cognitive decline in Alzheimer’s disease mice. J Exp Med 211, 1049–1062. (doi:10.1084/jem.20131751).

[100] Cortes-Canteli, M., Paul, J., Norris, E.H., Bronstein, R., Ahn, H.J., Zamolodchikov, D., Bhuvanendran, S., Fenz, K.M. & Strickland, S. 2010 Fibrinogen and beta-amyloid association alters thrombosis and fibrinolysis: a possible contributing factor to Alzheimer’s disease. Neuron 66, 695–709. (doi:10.1016/j.neuron.2010.05.014).

[101] Zamolodchikov, D. & Strickland, S. 2012 Abeta delays fibrin clot lysis by altering fibrin structure and attenuating plasminogen binding to fibrin. Blood 119, 3342–3351. (doi: 10.1182/blood-2011-11-389668).

[102] Zamolodchikov, D., Berk-Rauch, H.E., Oren, D.A., Stor, D.S., Singh, P.K., Kawasaki, M., Aso, K., Strickland, S. & Ahn, H.J. 2016 Biochemical and structural analysis of the interaction between beta-amyloid and fibrinogen. Blood 128, 1144–1151. (doi:10.1182/blood-2016-03-705228).

[103] Zamolodchikov, D. & Strickland, S. 2016 A possible new role for Abeta in vascular and inflammatory dysfunction in Alzheimer’s disease. Thromb Res 141 Suppl 2, S59–61. (doi: 10.1016/S0049-3848(16)30367-X).

